# Volitional Control of Individual Neurons in the Human Brain

**DOI:** 10.1101/2020.05.05.079038

**Authors:** Kramay Patel, Chaim N. Katz, Suneil K. Kalia, Milos R. Popovic, Taufik A. Valiante

**Affiliations:** Krembil Research Institute, Toronto Western Hospital (TWH), Ontario, M5T 1M8, Canada; Institute of Biomaterials and Biomedical Engineering, University of Toronto, Toronto, Ontario, M5S 3G9, Canada; Faculty of Medicine, University of Toronto, Toronto, ON, M5S 1A8, Canada; Division of Neurosurgery, Department of Surgery, University of Toronto, ON, M5S 1A1, Canada; The KITE Research Institute, University Health Network, Toronto, ON, M5G 2A2, Canada; Electrical and Computer Engineering, University of Toronto, Toronto, ON, M5S 3G4, Canada; CRANIA, University Health Network and University of Toronto, Toronto, ON, M5G 2A2, Canada

## Abstract

Can the human brain, a complex interconnected structure of over 80 billion neurons learn to control itself at the most elemental scale – a single neuron. We directly linked the firing rate of a single (direct) neuron to the position of a box on a screen, which participants tried to control. Remarkably, all subjects upregulated the firing rate of the direct neuron in memory structures of their brain. Learning was accompanied by improved performance over trials, simultaneous decorrelation of the direct neuron to local neurons, and direct neuron to beta frequency oscillation phase-locking. Such previously unexplored neuroprosthetic skill learning within memory related brain structures, and associated beta frequency phase-locking implicates the ventral striatum. Our demonstration that humans can volitionally control neuronal activity in mnemonic structures, may provide new ways of probing the function and plasticity of human memory without exogenous stimulation.

---

Advances in physical, and computational tools continue to inspire the development of devices to interrogate brain circuits and restore lost neural functioning. The motor system has long been a target for such devices, with an emerging interest in neuromodulatory as well as neuroprosthetic technologies for the interrogation and augmentation of cognition – in particular memory (*1–5*). The seminal work of Eberhard Fetz in the late 1960s, demonstrated that with the appropriate feedback and reward, monkeys can learn to control the activity of individual neurons in the primary motor cortex(*6, 7*). More recent work using advanced imaging and stimulation technologies in transgenic mice have demonstrated intentional neuroprosthetic learning of individual neurons within primary motor, and visual cortices (*8–12*). Whether such high fidelity neuroprosthetic skill learning is possible in the much larger and more architecturally complex human brain remains unknown. More specifically, it is unknown if such neuroprosthetic skills can be acquired in mnemonic structures that are not directly connected to the dorsal striatum (*13, 14*), a structure which appears essential for neuroprosthetic skill learning in the neocortex (*10–12*).

At a large spatial scales, scalp electroencephalography (EEG) has provided varied, albeit supportive literature regarding the efficacy of biofeedback to control oscillatory power in non-motor regions of the human brain (*15–19*). On a mesoscopic scale intracranial EEG (iEEG) recordings, have shown humans can control oscillations in the local field potential (LFP) within medial temporal lobe structures (*20, 21*). Few have even reported the possibility of controlling neuronal activity in medial temporal lobe (*22*), and other non-motor structures (*23*), however such control relied on invocation of previously identified concepts or motor imagery. Thus it remains unknown if operant conditioning of individual neurons within memory structures of the human brain is possible.

To explore this question we exploited the unique opportunity to obtain human single neuron recordings from epilepsy patients undergoing diagnostic iEEG to assess their surgical candidacy. We developed a closed-loop real-time instrumental learning task, where visual feedback is provided to participants as they try to learn to increase the firing rate of an arbitrarily chosen neuron. We show that: 1) humans can volitionally increase the firing rate of arbitrary individual neurons; 2) as with all forms of instrumental learning only a subset of participants get better at the task (learners); and 3) only learners demonstrated an increase in local spike field coherence (SFC), with the strongest SFC in the beta band, an uncommon oscillation observed in the human hippocampus. Our findings show that: 1) instrumental learning to control arbitrary individual neurons is possible in the human brain; 2) that such learning is possible outside of primary motor and sensory areas, and of particular interest in mnemonic structures, and; 3) intriguingly the unique beta band SFC signature of learners implicates the striatum, likely the ventral striatum, in instrumental learning within mnemonic structures (*14, 25*).

## Results

### Performing the Neurofeedback Task

We developed a neurofeedback task which required upregulation of the firing rate of an arbitrarily chosen neuron (henceforth called the direct neuron; Figure 1), from a mnemonic and/or non-motor brain region (Table S1. See Methods in Supplementary Materials for details on the choice of the direct neuron). Spiking activity of the direct neuron was sorted in real-time and convolved with a 200ms Gaussian (*21*) to obtain its smoothed instantaneous firing rate. The smoothed firing rate of the direct neuron was linearly mapped onto the vertical position of a square on a screen placed in front of the participant (see Methods in Supplementary Materials). Participants were instructed to try and move the block above a white horizontal line (threshold). Maintaining the box above threshold for over half a second indicated success. A successful trial was displayed, followed shortly by a distractor question, after which the next trial was triggered. In this way, it was ensured that each trial ended in a success. Testing was divided into blocks of 10 trials, and the participants were asked to finish the 10 trials in 10 minutes or less. To keep the participants motivated, we increased the difficulty of the next block of trials (by moving up the target line) if the previous 10 trials were completed in less than 5 minutes. Eleven participants completed a total of 17 sessions, where they controlled a different direct neuron in each session.

**Fig.1.**
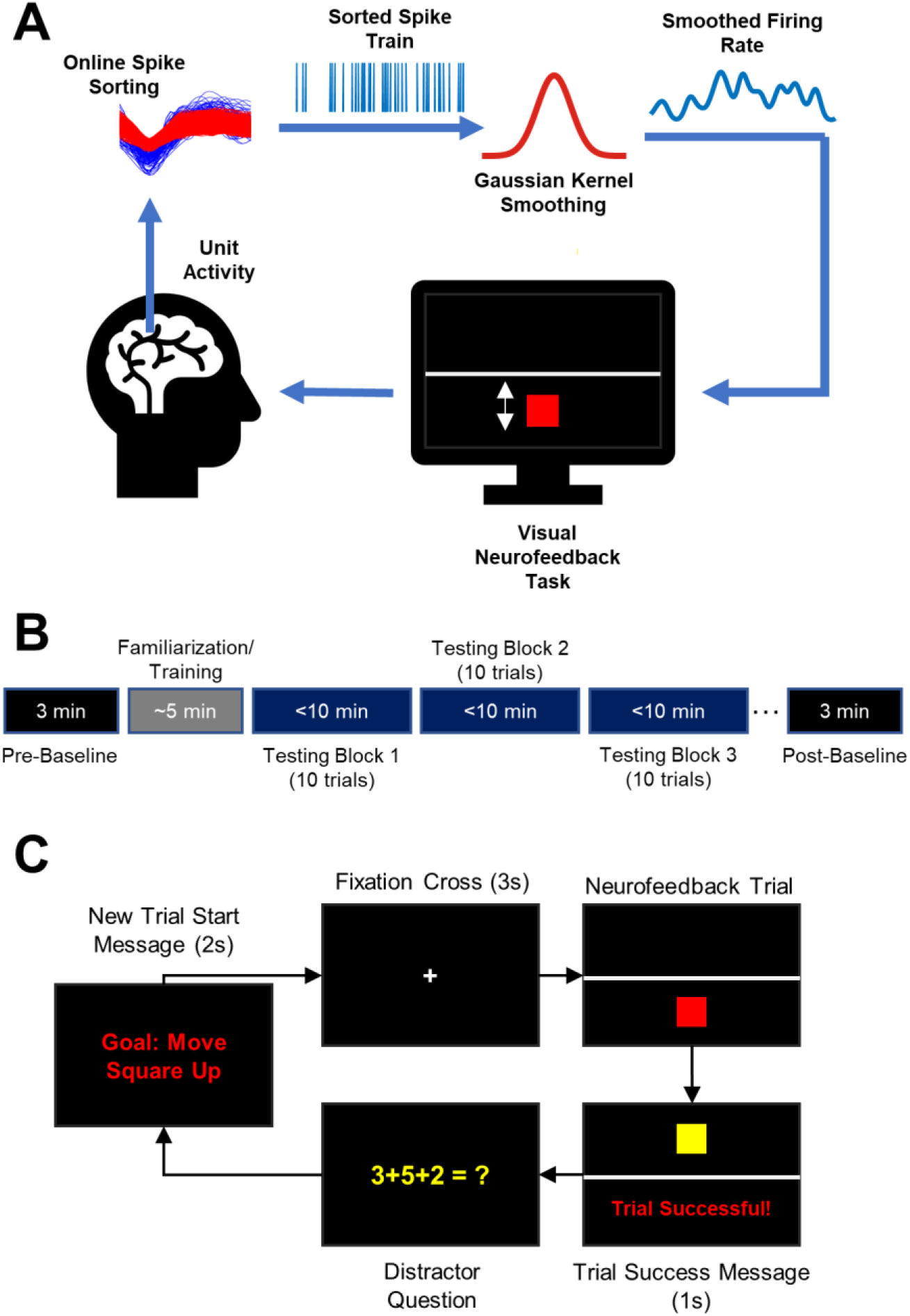
Visual neurofeedback task for modulating single neuron activity in the human brain. (A) Schematic showing the overall setup of the neurofeedback task. Single unit activity is extracted from the microwires in implanted Behnke Fried Macro Micro electrodes (AdTech) using the Neuralynx Atlas Digital Lynx system. Neurons from relevant channels were sorted online (using templates created with the KlustaKwik algorithm), and streamed over the Neuralynx NETCOM protocol to custom scripts in Matlab. These streamed, online spike trains were smoothed using a 200ms gaussian kernel to extract the instantaneous firing rate, which was then used as the control signal for the task. The neurofeedback task itself involves a red square moving vertically in response to the instantaneous firing rate of the chosen neuron. A horizontal white line indicates the target threshold, the crossing of which for 0.5 seconds results in a successful trial. (B) Schematic showing the experimental design, and order of a single testing session. An initial baseline session was performed, followed by 5 minutes of familiarization/training (during which an appropriate task difficulty is chosen). Following this, the testing phase began, which consists of at least 3 blocks (of 10 trials each). Each trial ended with a success, and each block was required to be finished in 10 minutes or less. Testing continued until a maximum of 12 blocks. Following testing, another baseline session was performed. (C) Schematic showing a single testing trial. Each trial began with a start message indicating the goal, followed by a fixation cross, followed by the actual trial. Each trial ended with a success message, followed by a quick distractor question involving the addition of 3 small integers.

All participants completed at least 30 trials (57±22 trials) indicating that all the participants were able to successfully upregulate the activity of their direct neuron (Figure 2A). Conversely, the firing rate of all other neurons recorded from the same bundle of microwires as the direct neuron (henceforth called indirect neurons) did not change prior to successful completion of the trial. Similarly, we did not observe a significant change in the firing rate of the 743 indirect neurons recorded from other microwire bundles prior to successful trial completion (Figure S1-A). To quantify this further we calculated the modulation depth of the direct neurons, defined as the average firing rate in the one second window after success subtracted from the average firing rate in the one second window before success. If success was triggered by random bursts of activity, the modulation depth would be close to zero since the bursts would likely continue into the post-success period. To the contrary, we saw a sharp decline in the activity of the direct neurons immediately following success (Figure 2B), resulting in a modulation depth significantly greater than zero (p<0.001, Single sample T-Test). To determine whether this type of upregulation was specific to the direct neuron, we calculated the modulation depth of indirect neurons. Indirect neurons’ firing rates were neither task-contingent, nor modulated with the direct neuron as evidenced by their modulation depth being close to zero (Figure 2B and Figure S1B).

**Fig. 2.**
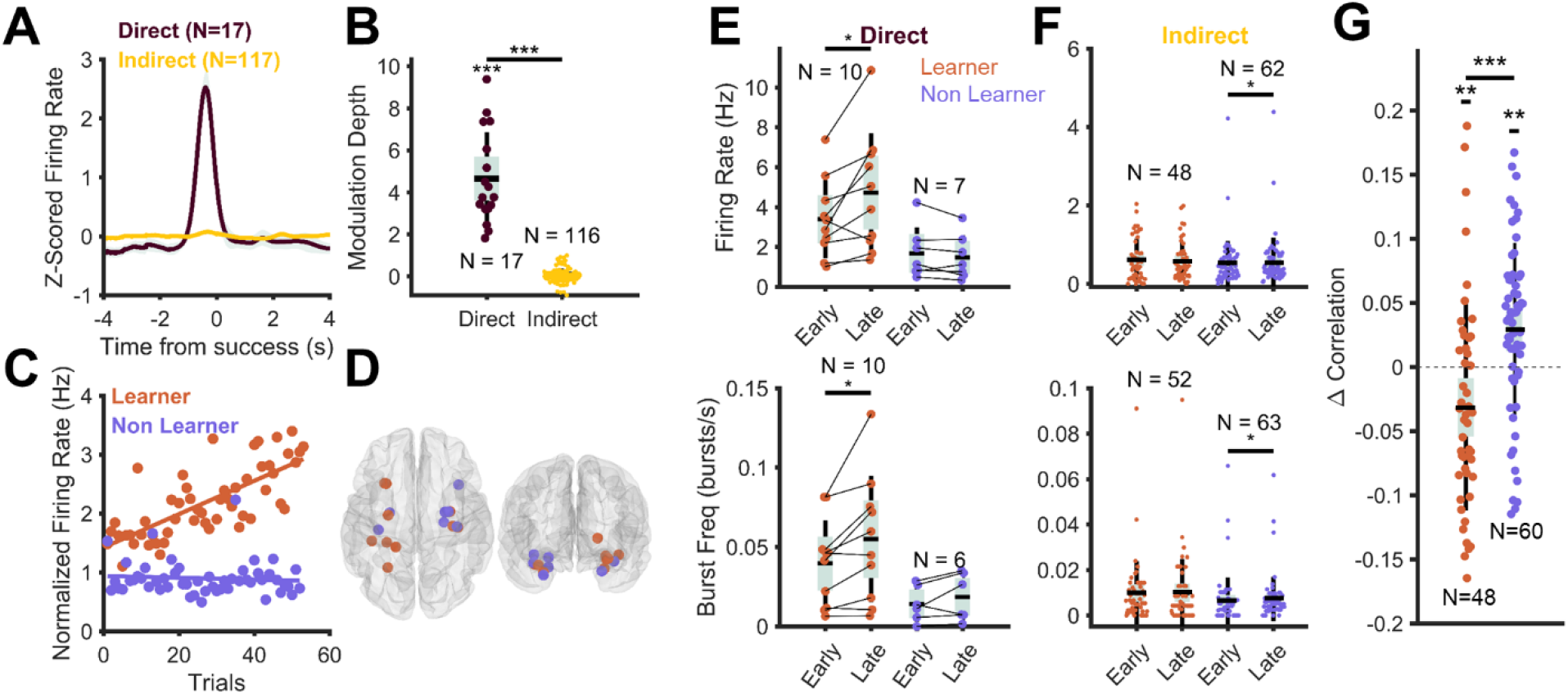
Learning to upregulate the activity of direct neurons using neurofeedback. (A) Firing rate of the direct neuron increased sharply immediately before success (t=0) peaking at 380ms before success, and returned to baseline immediately after success. Firing rate of indirect neurons was not modulated in the same epoch around success. (B) Modulation depth of direct neurons is significantly greater than zero (p<0.001, single sample two-tailed T Test) and significantly greater than that of indirect neurons (p<0.001, independent samples T Test). Outliers are removed using the Grubbs method. (C) Representative learner and non-learner sessions. A significant positive slope in the regression line between trial number and peak or average firing rate in each trial results was considered a learner session. All other sessions were defined as non-learner sessions. (D) Anatomical distribution of the direct neurons, color coded to match whether the session was a learner or non-learner. (E) Changes in the firing rate (top) and burst frequency (bottom) of direct neurons within a single session grouped by learner and non-learner sessions. Firing rate increased from the early to late trials in the learner sessions (p = 0.024, paired T-Test), and so does burst frequency (p = 0.019, paired T-Test). Changes in the firing rate (p = 0.21, paired T-Test) and burst frequency (p = 0.20, paired T-Test) are not evident for the non-learner sessions. (F) Same as in (E), but for indirect neurons. Firing rate and burst frequency do not change in learner sessions (Firing rate: p = 0.99, Wilcoxon Sign Rank Test; Burst Frequency: p = 0.98, paired T-Test), but increase significantly for the non-learner sessions (Firing rate: p = 0.018, Wilcoxon Sign Rank Test; Burst Frequency: p = 0.045, paired T-Test). (G) Change (late minus early) in spike correlations between the direct neuron and the neighbouring indirect neurons recorded from the same bundle of micro-wires. Correlations with neighbouring neurons decreased significantly in the learner sessions (p = 0.0094, single sample two-tailed T-Test), and increased significantly in the non-learner sessions (p = 0.0013, single sample two-tailed T-Test). Change in correlation is significantly different between the learner and non-learner sessions (p < 0.001, independent samples T-Test). For all figures, * indicates p<0.05, ** indicates p<0.01 and *** indicates p<0.001. For all figures, N indicates the number of neurons.

### Learning to Improve Performance

Neuroprosthetic skill learning is not uniformly acquired, where up to 30% of subjects are “non-learners” (*26*). We classified each session as a “learner” session when the participant was able to upregulate the average and/or peak firing rate of the direct neuron within the session (*20*). To do this, we performed a linear regression between the average and peak firing rate of the direct neuron as a function of the trial number. Sessions were defined as learner sessions if there was a significantly positive trend in either the peak or average firing rate of the direct neuron (Figure 2C; see Methods in Supplementary Materials for more details). With this definition, we defined 10 sessions as learners (across 7 patients) and the remaining 7 sessions as non-learners (see Supplementary Table 1 for participant demographics). Thus, while all participants were able to upregulate the activity of the direct neurons, only during some sessions were they able to improve their performance in the task.

As expected, the average firing rate of the direct neuron in learner sessions was significantly higher in the later trials (i.e. the last 15 trials) compared to the early trials (i.e. the first 15 trials) (Figure 2E, top panel). Similarly, the burst frequency of the direct neurons (calculated using a modified Poisson-surprise method) also increased significantly in the learner sessions but not in the non-learner sessions (Figure 2E, bottom panel). Indirect neurons demonstrated the opposite trend, with average firing rate and burst frequency increasing (by a small albeit significant magnitude) in the non-learner sessions, but not in the learner sessions (Figure 2F). Interestingly, the firing rate or burst frequency of indirect neurons recorded from other brain regions did not change from early to late trials in learners or non-learners (Figure S1-C and D). There is thus a stark dissociation between the neural activity between the direct and indirect neuronal populations, where during learner sessions, participants selectively modulate the activity of the direct neurons, and in non-learner sessions they unknowingly modulate the activity of neighbouring neurons, while failing to modulate the direct neuron. Thus, in the human brain, learning is accompanied by selective, volitional control over the direct neuron, whereas unsuccessful learning is characterized by non-specific modulation of the entire neural subpopulation consisting of both direct and indirect neurons. This dissociation is further exemplified by a decorrelation of the activity of the direct neuron from the neighbouring indirect neurons in the learner sessions, and an increase int his correlation in the non-learner sessions (Figure 2G). This finding mirrors similar findings reported using calcium imaging studies in rodents.(*9*).

### Spike-Field-Coherence develops During Learning

During neuroprosthetic skill acquisition in rodents, learning is accompanied by increased cortico-striatal communication evidenced by cortico-striatal coherence observed in the LFP (*11*), as well as spike field coherence (SFC) between cortical neurons and striatal oscillations and vice versa (*10–12*). Striatum is not a clinical target in iEEG recordings in epilepsy patients, and thus we were unable to test the hypothesis of striatal communication in the volitional control of individual neurons in humans. However we used rodent SFC findings to motivate a similar analysis to infer ‘communication through coherence’ (*27*) if such SFC was observed. Towards this end we computed the SFC between direct neurons and the LFP recorded by the closest macro contact of the Behnke-Fried electrode (local LFP; Figure 3A). For learners we found a striking increase in the SFC in the 10-20Hz range immediately before success (Figure 3A&B), while the non-learner population displayed no such increase in SFC. The ability to learn this skill is thus associated with a unique electrophysiological state of the brain (*28*), evidenced by increased SFC in the beta frequency range, and likely different from other “learning” states of the human brain (*29*).

**Fig. 3.**
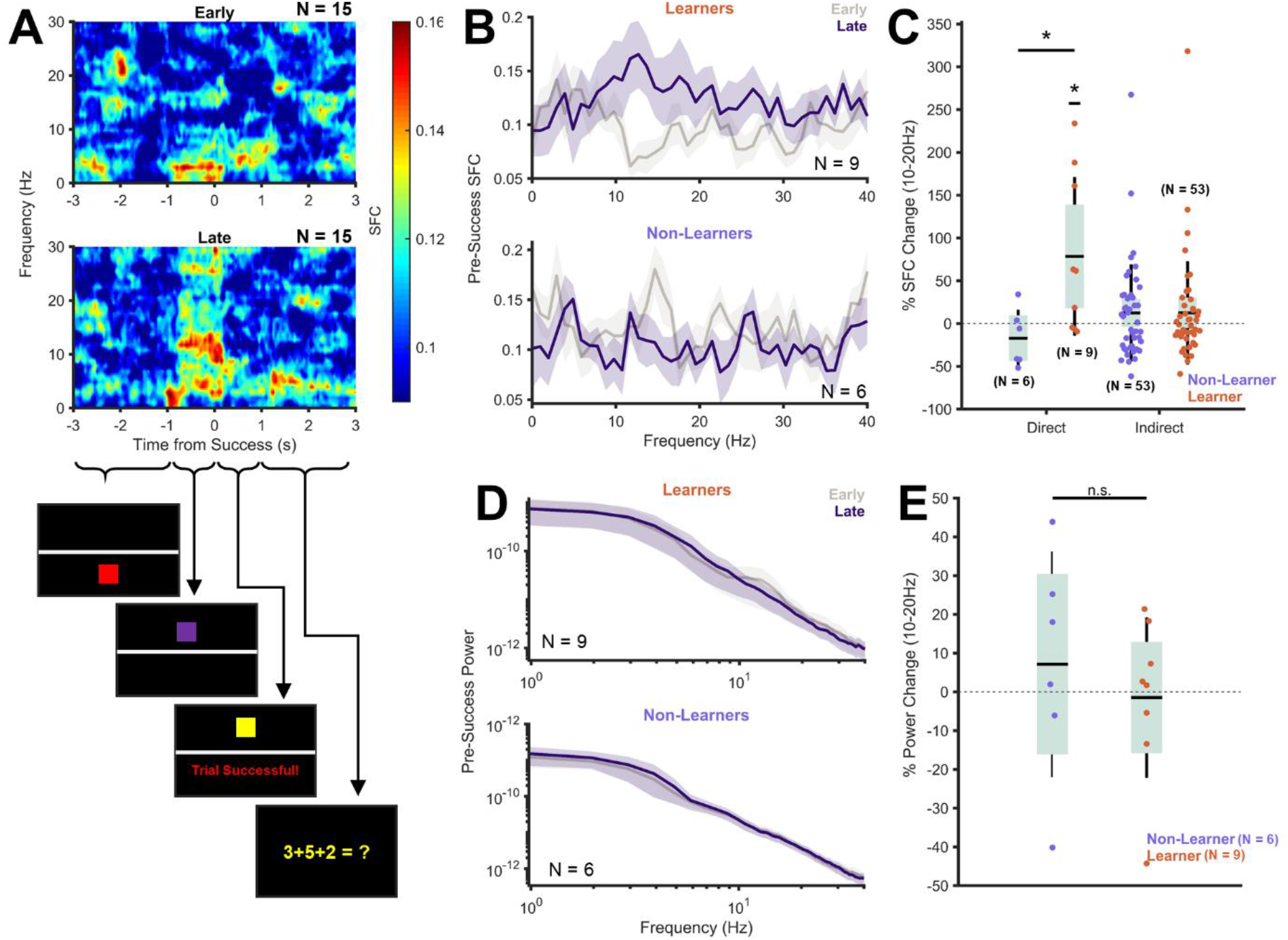
Local Spike-Field-Coherence in the 10-20Hz range emerges as learning progresses. (A) Coherogram of grand-average spike field coherence of the direct neurons in early (top) and late (bottom) trials. Schematics of the visual state of the task during each of the relevant periods (below). Note the significant increase in the spike field coherence (SFC) in the 10-20Hz band immediately preceding success (See Methods in Supplementary Material for details regarding the SFC calculation). (B) The grand average SFC in a 1 second window immediately preceding success. Notice the substantial increase in the SFC in the 10-20Hz range in the learner sessions, but not in the non-learner sessions. (C) Percent (%) change in SFC in the 10-20Hz range (from early to late trials) in the learner and non-learner sessions, for direct and indirect neurons. For the direct neurons, SFC increased significantly in the learner sessions (p = 0.035, single sample two-tailed T-Test) but not in the non-learner sessions (p = 0.26, single sample two-tailed T-Test). The % change in SFC was also significantly higher in learner sessions compared to non-learner sessions (p = 0.032, independent samples T-Test). For the indirect neurons, there is no change in the learner (p = 0.76, Wilcoxon Sign Rank Test) or the non-learner sessions (p = 0.37, Wilcoxon Sign Rank Test). (D) Grand average power spectra in the 1 second window immediately preceding success for learner (top) and non-learner (bottom) sessions. There were no significant learning related changes in the power spectrum in the pre-success interval, in the 10-20Hz frequency range. (E) Percentage power change in learner and non-learners is not different from 0 (p = 0.85 Learners, p = 0.57 Non-Learners, single sample two-tailed T-Test), or from each other (p = 0.53, independent samples T-Test). For all figures, * indicates p<0.05.

Since indirect neurons are not task relevant (their firing rates did not contribute to success), we anticipated that these indirect neurons would not develop the same learning-related SFC that was observed for direct neurons. To test this hypothesis, we calculated the SFC for indirect neurons to the local LFP and found no learning-related changes in both the learner and non-learner populations (Figure 3C). Prior to calculating the SFC, the firing rate of early and late trials were matched using a spike thinning procedure to prevent any biases resulting from unequal firing rates between the conditions (see Methods for details). Phase-related measures can often be affected by changes in oscillatory power (*30*). To determine whether the observed change in learning-related SFC was affected by spectral power changes, we computed the power spectra in the same time period in early and late trials. We found no differences in the power in the 10-20Hz frequency bands (Figure 3 D and E). Thus, in the absence of firing rate and power-related changes, the observed changes in the learning-related SFC must be driven by changes in spike timing immediately before success.

Additionally, we observed instances where the same participant could learn successfully in one session, but not in another (Supplementary Table 1). Despite this, we observed the learning related SFC changes confined to the learner sessions, suggesting the specificity of these changes to the act of learning itself, and not to other demographic factors.

The observed SFC in the beta band might be due to volume conducted low frequency oscillations. To address this specifically we calculated the SFC between the direct neurons and the LFP at non-local macro electrodes throughout the brain (Figure S2). In addition to the increase in the 10-20Hz frequency band SFC between the direct and non-local LFP was (Figure S2-D), an even more profound increase in theta frequency SFC observed in the SFC of the direct neuron to non-local LFP. Since theta is a ubiquitous oscillation in the human brain (*31*), including the human hippocampus (*29, 32*) and more likely to contribute to volume conduction (*33*), the beta frequency learning related SFC appears to be both specific in frequency range and local to the direct neuron during neuroprosthetic skill learning.

### Learning Related Spike-Field-Coherence Is Distinct from Anticipatory Reward

Since the participants were asked to hold the square above the threshold for more than 500ms, we wondered whether the observed SFC in the period immediately before success was driven by a reward anticipation mechanism (*34*). To test this theory, we extracted epochs around unsuccessful threshold crossings, i.e. points in time when the firing rate of the direct neuron crossed the threshold but for an insufficient time to trigger a successful trial. We hypothesized that if the 10-20Hz SFC we observed in the pre-success period was indeed the result of an anticipatory reward mechanism, we would observe a similar increase in the 10-20Hz SFC immediately after unsuccessful threshold crossings. Arguing against such an anticipatory reward mechanism, the spike-field-coherogram of the threshold crossing-aligned epochs (Figure 4A) did not demonstrate an increase in the 10-20Hz SFC immediately after threshold crossings (Figure 4B, top panel). Furthermore, there was no change in the SFC in this frequency band in the post-threshold crossing window between the early and late trials (Figure 4C) for learners and non-learners, confirming that this type of reward anticipation does not drive the learning related SFC changes observed in the success aligned epochs.

**Fig. 4.**
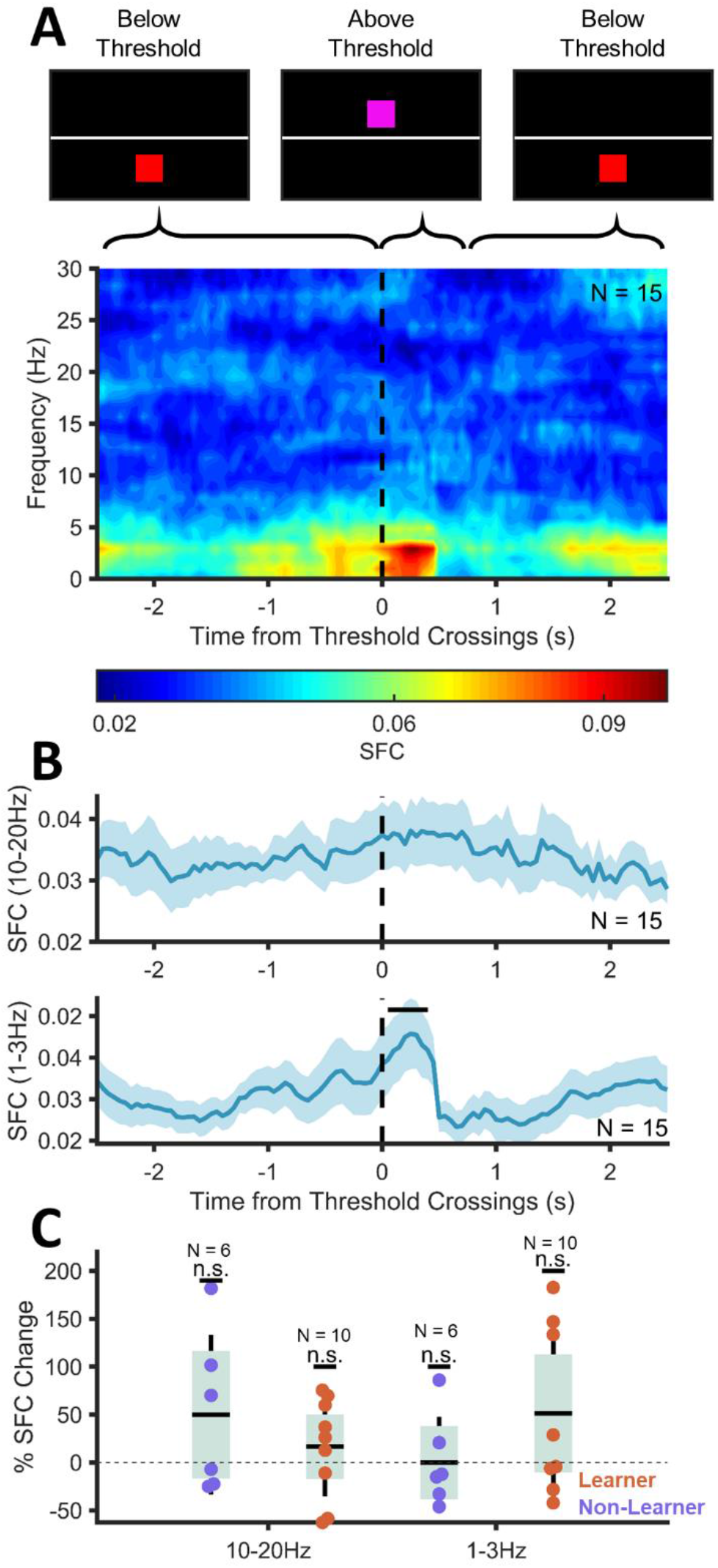
Unrewarded threshold crossings do not result in a change in SFC in the 10-20Hz band. (A) Spectrogram showing the Spike-Field Coherence between the direct neurons and the local LFP aligned to unrewarded threshold crossings. Notice that there is no significant increase in SFC in the 10-20Hz range following the threshold crossings. (B) Grand average SFC in the 10-20Hz band (top) and 1-3Hz band (bottom). Note that SFC does not increase in the 10-20Hz band following threshold crossings, but it does increase significantly following threshold crossings in the 1-3Hz band (significant portions indicated with a bold line on top of the graph, p<0.05 non-parametric permutation testing with random time-shuffles, 2000 iterations). (C) Change in SFC between early and late trials in the 10-20Hz and 1-3Hz frequency ranges for learners and non-learners. No significant changes observed (10-20Hz: Learner – p = 0.37, Non-Learner p = 0.20 single sample two-tailed T Test; 1-3Hz: Learner – p = 0.45, Non-learner – p = 0.99, single sample two-tailed T-Test). No learning-related changes in SFC observed in either frequency band following threshold crossing.

Interestingly, we did observe a significant increase in the delta-band (1-3Hz) SFC immediately following threshold crossings (Figure 4B). This finding was concordant with the increased delta SFC observed in the window immediately surrounding success (Figure 3A). To determine whether this delta SFC was learning related, we compared the SFC in the post-threshold crossing window in the early vs the late trials (Figure 4C). We observed no significant difference in the delta SFC in this window in the early vs. late trials (for learners and non-learners), suggesting that the delta-SFC was not learning-related, and likely related to the design of the neurofeedback task. Consistent with this hypothesis the delta-SFC increase was associated with a delta power increase in a similar time window (Figure S3). This suggests that the observed delta-SFC increase is likely driven by the image onset evoked response due to the colour change of the square from red to purple when it crosses the threshold (*32*) rather than either anticipatory reward, or a learning related mechanism.

## Discussion

Here we demonstrate, using a visual neurofeedback task, that humans can learn to upregulate the activity of arbitrarily chosen neurons in their brain in a highly specific and volitional manner. Our results compliment non-human primate and rodent single neuron neuroprosthetic skill learning research and extend the possibility of such learning beyond previous work in sensorimotor cortices to associational structures of the brain, in particular the limbic system.

A large body of existing literature provides evidence for this type of neuroprosthetic skill learning in the motor cortex of rodents (*9*–*12*, *35*), and primates (*6*, *7*, *36*–*38*). Control at the single neuron level in the motor cortex has been shown to require the dorsal striatum (*11, 12*), which serves as an input tier for the basal ganglia. Hence, neuroprosthetic skill learning in the motor cortex is largely analogous to motor learning, in which cortico-basal ganglia loops facilitate an action selection process where competing motor programs are either inhibited or released from inhibition. This is facilitated by parallel direct and indirect pathways which allow disinhibition and inhibition, respectively, of neuronal ensembles in the sensorimotor cortices, allowing for selection of a contextually relevant motor program (*39*). Similar cortico-basal ganglia loops are implicated in selection and generation of a variety of different cognitive patterns which may facilitate more abstract skill learning (*40*). In fact, recent rodent studies provide conclusive evidence that animals can modulate highly specific neuronal activity in primary sensory cortex which again is dependent on the dorsal striatum, similar to learning in the motor cortex (*10*). Since the majority of the neocortex projects to the dorsal striatum (*13, 41*), we anticipate that this type of neuroprosthetic skill learning may be possible in most of the neocortex.

In this study however, we demonstrate that this type of learning is also possible in the paleo-cortex of the human brain, as well as other non-motor, non-sensory regions. These structures are largely dissociated from the dorsal striatal system (*14*). However, despite this dissociation, we demonstrated that participants learned to modulate activity in a specific and volitional manner, much like other neocortical regions explored in non-human primates and rodents. Motor skill learning, and neuroprosthetic skill learning, proceeds in a prototypical manner, where the early phase of learning is characterized by a rapid acquisition of task parameters, following by a slower refinement process (*10*–*12*, *28*). The experimental sessions in this study were not long enough to investigate the later stages of learning, but we robustly demonstrate the early stage of learning, characterized by rapid changes in the firing characteristics of the direct neurons. While limbic structures do not directly project to the dorsal striatum, they do project heavily to the ventral striatum. In fact, the ventral striatum is thought to serve as the interface between the limbic and motor systems (*25, 42*). So is it possible that the type of neuroprosthetic skill learning that we demonstrate here is facilitated by the ventral striatum instead?

While we cannot answer this question by directly recording activity from the ventral striatum in humans, we sought out signatures of this interaction, where we observed an increase in SFC in the high alpha/low beta bands as learning progressed. This increase in SFC was independent of power or firing rate changes, was specific to the direct neurons, and occurred only in learning sessions. In the medial temporal lobe, oscillations in this frequency band are rarely observed and in particular have not been reported in human MTL regions where delta, theta, and gamma frequency activity has been associated with a myriad of behaviors (*32*, *43*–*45*) including neuroprosthetic skill learning (*20, 21*). In rodents Lansink and colleagues demonstrated beta oscillations in the hippocampus driven by reward predictive cues, and enhanced by learning (*34*). They also demonstrated the presence of hippocampal spiking activity phase-locked to the underlying beta oscillations and driven by reward-predictive cues. Interestingly, they also observed increased SFC coherence between neurons in the ventral striatum and beta oscillations in the hippocampus in response to reward predictive cues. Thus, beta oscillations in the hippocampus and related structures may be driven by a reward prediction mechanism, potentially driven by the ventral tegmental afferents to CA1 (*46*), or indirectly from the striatum via the ventral pallidal-mediodorsal thalamic route (*47*). When we aligned our data to unsuccessful reward crossings, we did not observe any reward-predictive increases in beta power or synchrony (Figure 4). Furthermore, Lansink and colleauges reported concomitant reward-predictive increase in theta power and theta SFC, which were also not present in our findings. Thus, the rodent studies suggest to us that the SFC that we observe in the beta frequency range: 1) may indeed reflect MTL-ventral striatal communication through coherence; 2) is unlikely to be a reward prediction signal, and; 3) is akin to the cortico-striatal coherence seen during instrumental learning in rodents and non-human primates (NHP).

Unlike limbic structures, beta oscillations are commonplace in the striatal system, in rodents (*25, 48*) and NHPs (*49, 50*), including the ventral striatum of rodents (Berke, 2009; Berke et al., 2004) and the nucleus accumbens in humans (*51*). In rodents Neely and colleagues investigated neuroprosthetic skill learning in the primary visual cortex (V1), while simultaneously recording from the dorsomedial striatum (which receives direct projections from the primary visual cortex) (*41, 52*). They demonstrated that as animals learned to control specific neuronal activity in V1, they increasingly recruited the striatum, demonstrated by an increased spiking of striatal neurons along with concomitant increases in beta and gamma power (*10*). Furthermore, and perhaps most intriguingly, they demonstrated that as learning progressed, spiking activity of the direct units in V1 became more coherent with the local field potentials in the 10-25Hz band. This finding mirrors our results in the human limbic structures, although we did not observe changes in oscillatory power in this frequency band as they reported. Since attention has actually been shown decrease alpha/beta band SFC in the visual cortex (*53*), the increased SFC we observe in the alpha/beta SFC is unlikely due to attentional modulation. Similarly, Koralek and colleagues demonstrated that neuroprosthetic skill learning in the primary motor cortex (M1) is accompanied by increased success-aligned spike field coherence between M1 spikes and striatal LFP in the 6-14Hz band (*11*). In light of these results our SFC findings likely reflect the signature of increased communication between MTL-ventral striatal ensembles that underlie the learning of the visual neuroprosthetic skill.

One of the canonical characteristics of the cortico-basal ganglia loops is the presence of parallel inhibitory and disinhibitory pathways (*39, 54*), which allow the basal ganglia to play a role in selection of context relevant motor plans or even cognitive strategies (*40*). The medium spiny neurons (MSNs), which are ubiquitous within the basal ganglia, are furnished with dopamine receptors in close proximity to the corticostriatal terminals (*55*). Dopaminergic innervation of these MSNs by the midbrain dopaminergic system facilitates plastic synaptic changes which shapes striatal, and the resulting basal ganglia outputs, playing a role in facilitating reward-based learning. While the dorsal striatum is unlikely to be involved in the limbic neuroprosthetic skill learning demonstrated here, the ventral striatum is also known to form cortico-basal ganglia loops (*54*), with a variety of limbic structures and the anterior cingulate cortex (ACC) as its primary input and output (*14*, *56*–*58*). Since the MSNs that compose much of the striatum are difficult to excite (*39*), convergent input from the limbic structures and the ACC could drive ventral striatal MSNs, activating a series of parallel inhibitory and disinhibitory circuits that can be actively tuned via the midbrain dopaminergic system to facilitate reward-based learning of precise limbic activity patterns. Future work in animal models will certainly focus on interrogating this limbic-basal ganglia circuitry to establish the significance of the ventral striatum in facilitating this type of limbic neuroprosthetic skill learning.

The data presented here suggest that single neuron activity in limbic structures can be precisely regulated in a rapid, highly specific and volitional manner in humans. Furthermore, this type of neuroprosthetic skill learning in limbic structures is likely facilitated by the limbic-basal ganglia circuity involving the ventral striatum. Such, high fidelity self-regulation of neural activity may provide an avenue for the development of novel neuroprosthetics for the treatment of neurological conditions which commonly present with pathological activity in limbic structures, such as medically refractory epilepsy. Furthermore since limbic structures, and particularly those of the medial temporal lobe, are critical to mnemonic processes, obtaining volitional control over highly specific activity in these structures may provide a mechanism of probing the function and plasticity of these brain structures without exogenous stimulation.

## Supporting information

Supplementary Items

## Acknowledgements

We would like to acknowledge Victoria Barkley for her help with organizing and assisting with patient data collection, Andrea Gómez Palacio for her help with patient data collection, Ryan Ramos for his contributions towards developing the neurofeedback task, and David Groppe for his advice on choosing appropriate statistical analyses.

## Author Contributions

Conceptualization – K.P., T.V.; Data curation – K.P; Formal Analysis – K.P.; Funding Acquisition – T.V.; Investigation – K.P., C.K.; Methodology – K.P.; Resources – K.P., T.V., S.K.; Software – K.P.; Supervision – M.P, T.V.; Visualization – K.P. Writing (original draft) – K.P.; Writing (review and editing) – K.P., C.K., M.P., S.K., T.V.

## Data and Materials Availability

The spike detection and sorting toolbox used for offline sorting (OSort), and the Chronux toolbox (use for spectral analysis) are both available as an open source toolboxes. Data and custom MATLAB scripts used for analysis here are available upon request from Taufik A. Valiante.

---

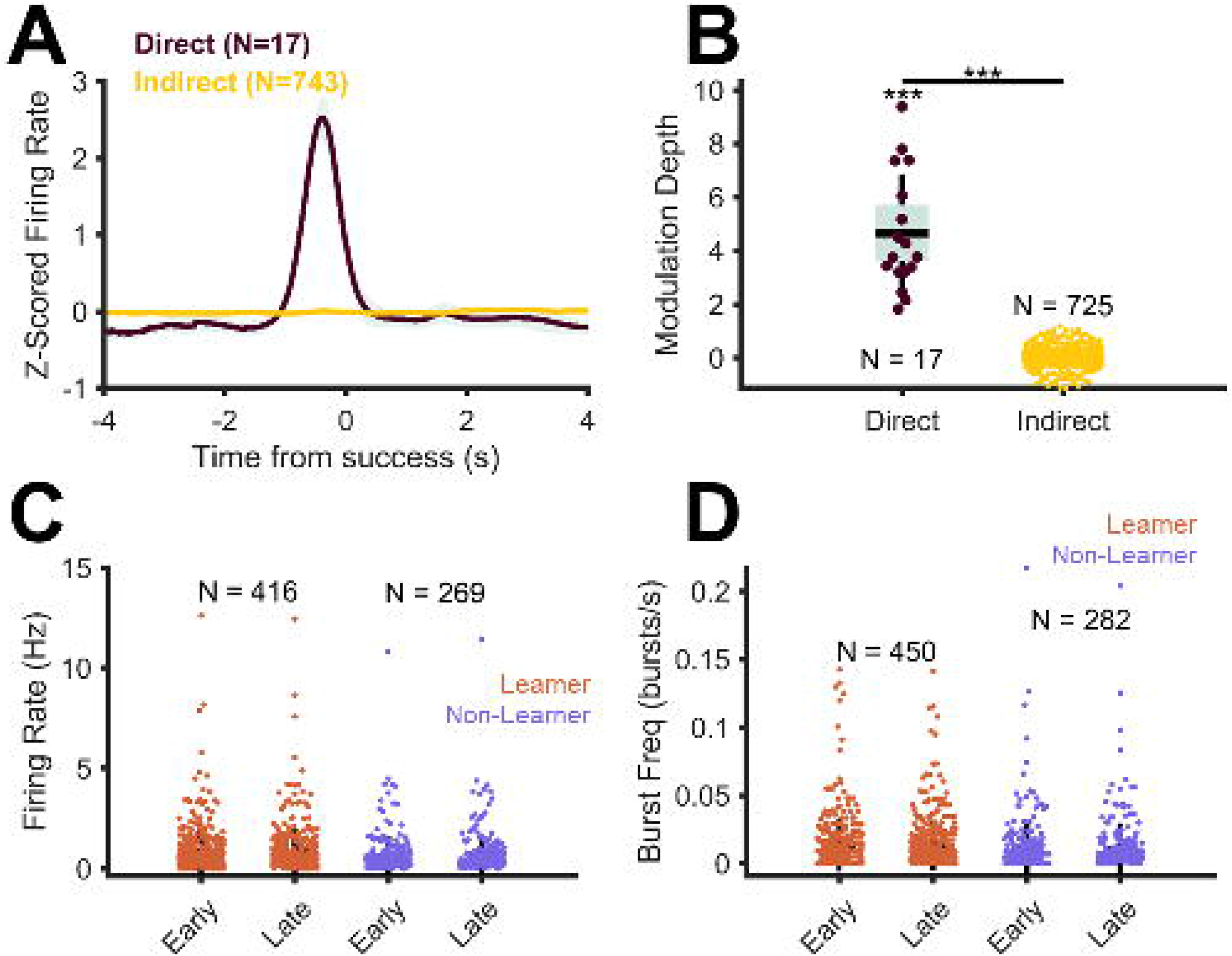

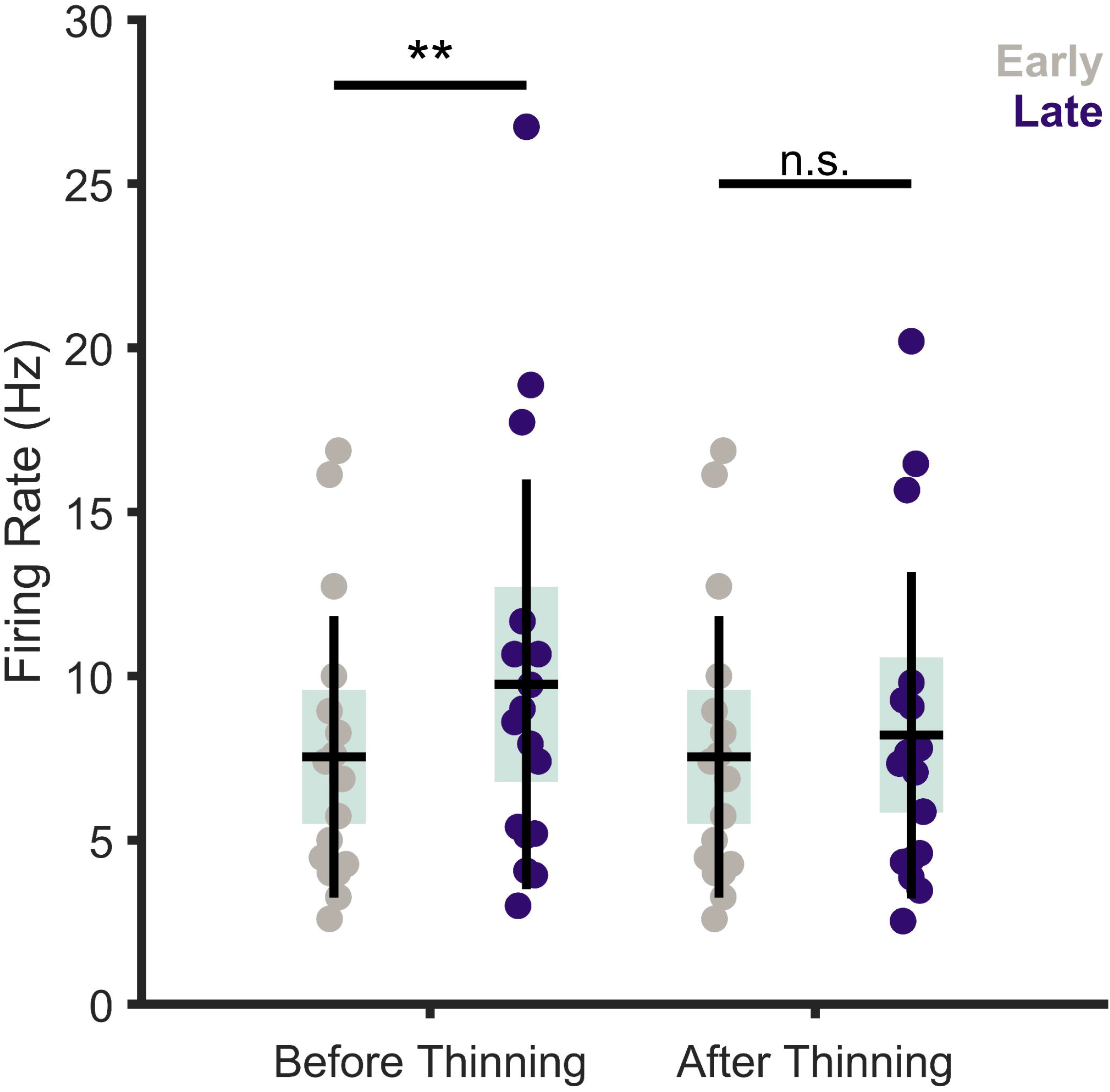

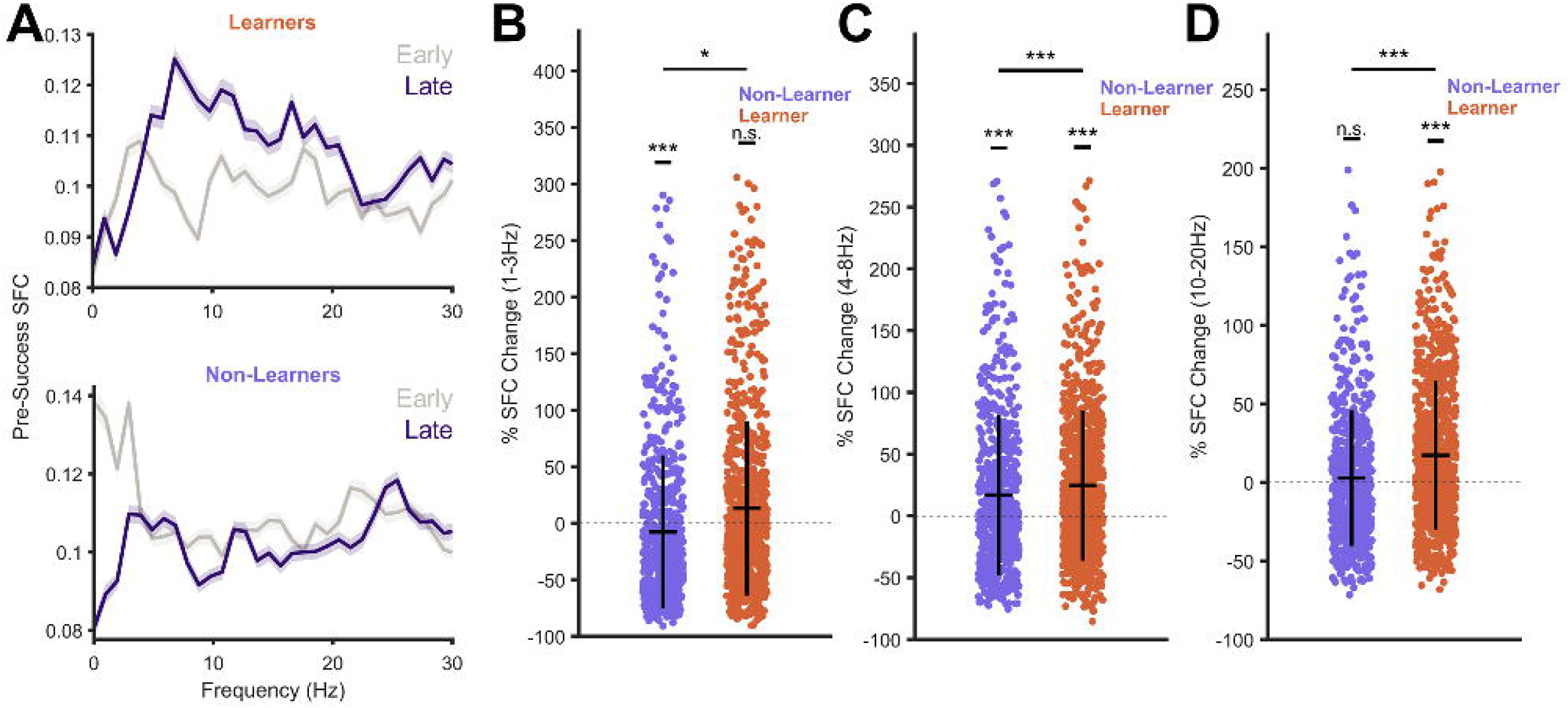

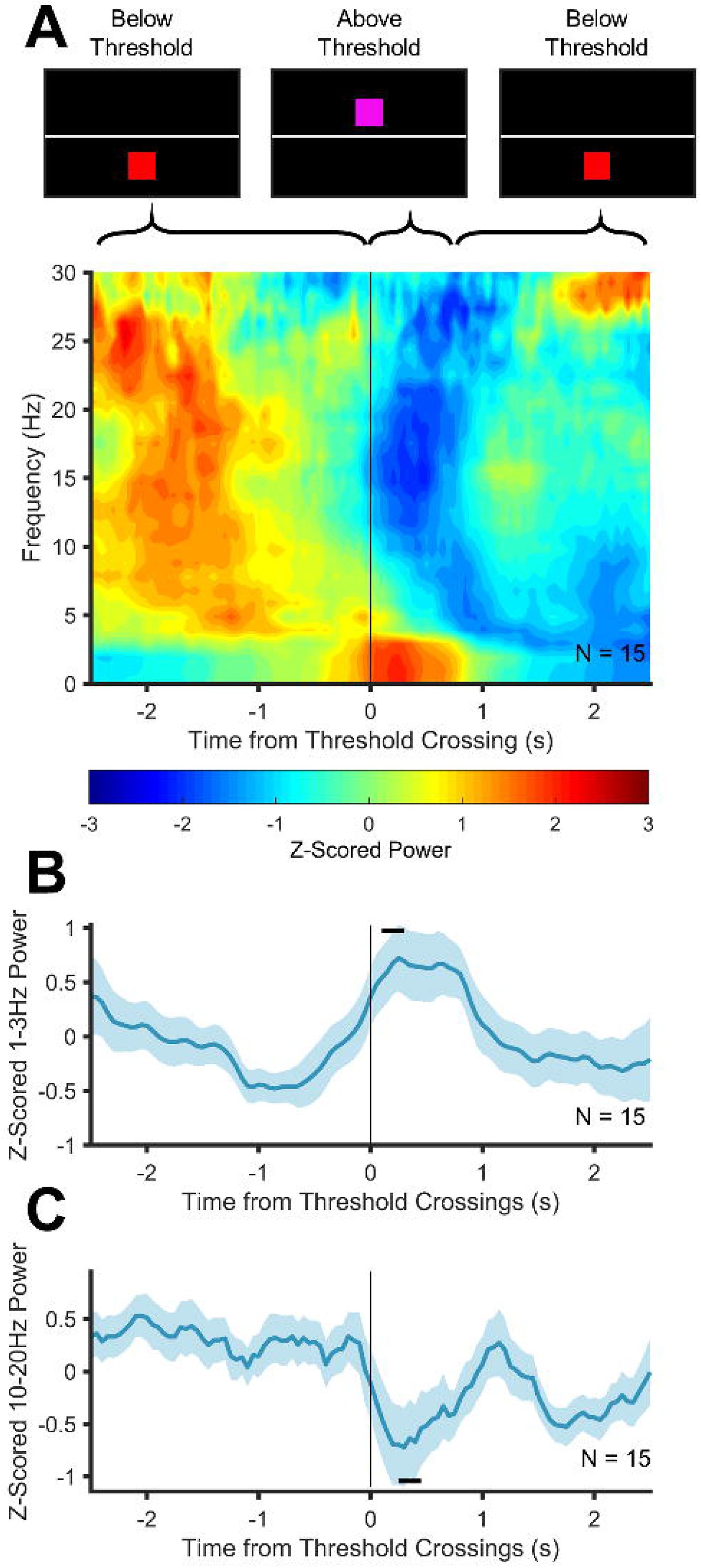

## References

1. A. M. Lozano, L. Fosdick, M. M. Chakravarty, J. M. Leoutsakos, C. Munro, E. Oh, K. E. Drake, C. H. Lyman, P. B. Rosenberg, W. S. Anderson, D. F. Tang-Wai, J. C. Pendergrass, S. Salloway, W. F. Asaad, F. A. Ponce, A. Burke, M. Sabbagh, D. A. Wolk, G. Baltuch, M. S. Okun, K. D. Foote, M. P. McAndrews, P. Giacobbe, S. D. Targum, C. G. Lyketsos, G. S. Smith, A Phase II Study of Fornix Deep Brain Stimulation in Mild Alzheimer’s Disease. J. Alzheimer’s Dis. 54, 777–787 (2016).

2. D. J. Lee, A. M. Lozano, Current Status of Deep Brain Stimulation for Alzheimer’s Disease: From Chance Observation to Clinical Trials. Cold Spring Harb. Symp. Quant. Biol. LXXXIII, 037440 (2019).

3. J. Kuhn, K. Hardenacke, D. Lenartz, T. Gruendler, M. Ullsperger, C. Bartsch, J. K. Mai, K. Zilles, A. Bauer, A. Matusch, R. J. Schulz, M. Noreik, C. P. Bührle, D. Maintz, C. Woopen, P. Häussermann, M. Hellmich, J. Klosterkötter, J. Wiltfang, M. Maarouf, H. J. Freund, V. Sturm, Deep brain stimulation of the nucleus basalis of Meynert in Alzheimer’s dementia. Mol. Psychiatry. 20, 353–360 (2015).

4. S. Hescham, L. W. Lim, A. Jahanshahi, H. W. M. Steinbusch, J. Prickaerts, A. Blokland, Y. Temel, Deep brain stimulation of the forniceal area enhances memory functions in experimental dementia: The role of stimulation parameters. Brain Stimul. 6, 72–77 (2013).

5. Y. Ezzyat, P. A. Wanda, D. F. Levy, A. Kadel, A. Aka, I. Pedisich, M. R. Sperling, A. D. Sharan, B. C. Lega, A. Burks, R. E. Gross, C. S. Inman, B. C. Jobst, M. A. Gorenstein, K. Davis, G. A. Worrell, M. T. Kucewicz, J. M. Stein, R. Gorniak, S. R. Das, D. S. Rizzuto, M. J. Kahana, Closed-loop stimulation of temporal cortex rescues functional networks and improves memory. Nat. Commun. 9(2018), doi:10.1038/s41467-017-02753-0.

6. E. E. Fetz, M. A. Baker, Operantly conditioned patterns on precentral unit activity and correlated responses in adjacent cells and contralateral muscles. J. Neurophysiol. 36, 179–204 (1973).

7. E. E. Fetz, Operant conditioning of cortical unit activity. Science. 163, 955–8 (1969).

8. M. Prsa, G. L. Galiñanes, D. Huber, Rapid Integration of Artificial Sensory Feedback during Operant Conditioning of Motor Cortex Neurons. Neuron. 93, 929–939.e6 (2017).

9. K. B. Clancy, A. C. Koralek, R. M. Costa, D. E. Feldman, J. M. Carmena, Volitional modulation of optically recorded calcium signals during neuroprosthetic learning. Nat. Neurosci. 17, 807–809 (2014).

10. R. M. Neely, A. C. Koralek, V. R. Athalye, R. M. Costa, J. M. Carmena, Volitional Modulation of Primary Visual Cortex Activity Requires the Basal Ganglia. Neuron. 97, 1356–1368.e4 (2018).

11. A. C. Koralek, R. M. Costa, J. M. Carmena, Temporally Precise Cell-Specific Coherence Develops in Corticostriatal Networks during Learning. Neuron. 79, 865–872 (2013).

12. A. C. Koralek, X. Jin, J. D. Long, R. M. Costa, J. M. Carmena, J. D. Long II, R. M. Costa, J. M. Carmena, Corticostriatal plasticity is necessary for learning intentional neuroprosthetic skills. Nature. 483, 331–335 (2012).

13. J. M. Kemp, T. P. S. Powell, The cortico-striate projection in the monkey. Brain. 93, 525–546 (1970).

14. P. Voorn, L. J. M. J. Vanderschuren, H. J. Groenewegen, T. W. Robbins, C. M. Pennartz, Putting a spin on the dorsal-ventral divide of the striatum. Trends Neurosci. 27, 468–474 (2004).

15. J. F. Lubar, W. W. Bahler, Behavioral management of epileptic seizures following EEG biofeedback training of the sensorimotor rhythm. Biofeedback Self. Regul. 1, 77–104 (1976).

16. J. F. Lubar, H. S. Shabsin, S. E. Natelson, G. S. Holder, S. F. Whitsett, W. E. Pamplin, D. I. Krulikowski, EEG Operant Conditioning in Intractable Epileptics. Arch. Neurol. 38, 700–704 (1981).

17. T. Ros, B. J. Baars, R. A. Lanius, P. Vuilleumier, Tuning pathological brain oscillations with neurofeedback: a systems neuroscience framework. Front. Hum. Neurosci. 8(2014), doi:10.3389/fnhum.2014.01008.

18. R. Sitaram, T. Ros, L. Stoeckel, S. Haller, F. Scharnowski, J. Lewis-Peacock, N. Weiskopf, M. L. Blefari, M. Rana, E. Oblak, N. Birbaumer, J. Sulzer, Closed-loop brain training: The science of neurofeedback. Nat. Rev. Neurosci. 18(2017), pp. 86–100.

19. J. E. Walker, Using QEEG-guided neurofeedback for epilepsy versus standardized protocols: Enhanced effectiveness? Appl. Psychophysiol. Biofeedback. 35, 29–30 (2010).

20. J. Corlier, D. Rimsky-Robert, M. Valderrama, K. Lehongre, C. Adam, S. Clémenceau, S. Charpier, J. Bastin, P. Kahane, J. P. Lachaux, V. Navarro, M. Le Van Quyen, Self-induced intracerebral gamma oscillations in the human cortex. Brain. 139, 3084–3091 (2016).

21. J. Corlier, M. Valderrama, M. Navarrete, K. Lehongre, D. Hasboun, C. Adam, H. Belaid, S. Clémenceau, M. Baulac, S. Charpier, V. Navarro, M. L. Van Quyen, Voluntary control of intracortical oscillations for reconfiguration of network activity. Sci. Rep. 6(2016), doi:10.1038/srep36255.

22. M. Cerf, N. Thiruvengadam, F. Mormann, A. Kraskov, R. Q. Quiroga, C. Koch, I. Fried, On-line, voluntary control of human temporal lobe neurons. Nature. 467, 1104–1108 (2010).

23. 1 Tyson Aflalo, 1 * Spencer Kellis, 1 * Christian Klaes, 1 Brian Lee, 2 Ying Shi, 1 Kelsie Pejsa, 2 Kathleen Shanfield, 3 Stephanie Hayes-Jackson, 3 Mindy Aisen, 3 Christi Heck, 2 Richard A. Andersen, 1 Charles Liu, †, Decoding motor imagery from the posterior parietal cortex of a tetraplegic human. Science (80-.). 348, 906–910 (2015).

24. R. Q. Quiroga, Concept cells: the building blocks of declarative memory functions. Nat. Rev. Neurosci. 13, 587–597 (2012).

25. J. D. Berke, M. Okatan, J. Skurski, H. B. Eichenbaum, Oscillatory entrainment of striatal neurons in freely moving rats. Neuron. 43, 883–896 (2004).

26. E. M. Hammer, S. Halder, B. Blankertz, C. Sannelli, T. Dickhaus, S. Kleih, K.-R. Müller, Kübler, Psychological predictors of SMR-BCI performance. Biol. Psychol. 89, 80–86 (2012).

27. P. Fries, Rhythms for Cognition: Communication through Coherence. Neuron. 88, 220–235 (2015).

28. V. R. Athalye, J. M. Carmena, R. M. Costa, Neural reinforcement: re-entering and refining neural dynamics leading to desirable outcomes. Curr. Opin. Neurobiol. 60(2020), pp. 145–154.

29. U. Rutishauser, I. B. Ross, A. N. Mamelak, E. M. Schuman, Human memory strength is predicted by theta-frequency phase-locking of single neurons. Nature. 464, 903–907 (2010).

30. M. X. Cohen, Analyzing Neural Time Series Data (2014).

31. D. M. Groppe, S. Bickel, C. J. Keller, S. K. Jain, S. T. Hwang, C. Harden, A. D. Mehta, Dominant frequencies of resting human brain activity as measured by the electrocorticogram. Neuroimage. 79, 223–233 (2013).

32. C. N. Katz, K. Patel, O. Talakoub, D. Groppe, K. Hoffman, T. A. Valiante, Differential generation of saccade, fixation and image onset event-related potentials in the human mesial temporal lobe. bioRxiv, 442855 (2018).

33. L. Lalla, P. E. Rueda Orozco, M. T. Jurado-Parras, A. Brovelli, D. Robbe, Local or not local: Investigating the nature of striatal theta oscillations in behaving rats. eNeuro. 4, 128–145 (2017).

34. C. S. Lansink, G. T. Meijer, J. V Lankelma, M. A. Vinck, J. C. Jackson, C. M. A. Pennartz, Reward expectancy strengthens CA1 theta and beta band synchronization and hippocampal-ventral striatal coupling. J. Neurosci. 36, 10598–10610 (2016).

35. P. J. Arduin, Y. Frégnac, D. E. Shulz, V. Ego-Stengel, Bidirectional control of a one-dimensional robotic actuator by operant conditioning of a single unit in rat motor cortex. Front. Neurosci. (2014), doi:10.3389/fnins.2014.00206.

36. R. W. Eaton, T. Libey, E. E. Fetz, Operant conditioning of neural activity in freely behaving monkeys with intracranial reinforcement. J. Neurophysiol. 117, 1112–1125 (2017).

37. S. Kobayashi, W. Schultz, M. Sakagami, Operant conditioning of primate prefrontal neurons. J. Neurophysiol. 103, 1843–1855 (2010).

38. C. T. Moritz, E. E. Fetz, Volitional control of single cortical neurons in a brain-machine interface. J. Neural Eng. 8(2011), doi:10.1088/1741-2560/8/2/025017.

39. H. J. Groenewegen, The basal ganglia and motor control. Neural Plast. 10, 107–20 (2003).

40. A. M. Graybiel, The basal ganglia and cognitive pattern generators. Schizophr. Bull. 23, 459–469 (1997).

41. A. J. McGeorge, R. L. M. Faull, The organization of the projection from the cerebral cortex to the striatum in the rat. Neuroscience. 29, 503–537 (1989).

42. G. J. Mogenson, D. L. Jones, C. Y. Yim, From motivation to action: Functional interface between the limbic system and the motor system. Prog. Neurobiol. 14(1980), pp. 69–97.

43. M. J. Jutras, P. Fries, E. A. Buffalo, Oscillatory activity in the monkey hippocampus during visual exploration and memory formation. Proc. Natl. Acad. Sci. 110, 13144–13149 (2013).

44. M. J. Jutras, P. Fries, E. A. Buffalo, Behavioral/Systems/Cognitive Gamma-Band Synchronization in the Macaque Hippocampus and Memory Formation (2009), doi:10.1523/JNEUROSCI.0640-09.2009.

45. R. Montefusco-Siegmund, T. K. Leonard, K. L. Hoffman, Hippocampal gamma-band Synchrony and pupillary responses index memory during visual search. Hippocampus. 27, 425–434 (2017).

46. H. J. Groenewegen, H. W. Berendse, S. N. Haber, Organization of the output of the ventral striatopallidal system in the rat: Ventral pallidal efferents. Neuroscience. 57, 113–142 (1993).

47. H. J. Groenewegen, C. I. Wright, H. B. M. Uylings, The anatomical relationships of the prefrontal cortex with limbic structures and the basal ganglia. J. Psychopharmacol. 11, 99–106 (1997).

48. D. K. Leventhal, G. J. Gage, R. Schmidt, J. R. Pettibone, A. C. Case, J. D. Berke, Basal ganglia beta oscillations accompany cue utilization. Neuron. 73, 523–536 (2012).

49. H. Merchant, R. Bartolo, Primate beta oscillations and rhythmic behaviors. J. Neural Transm. 125(2018), pp. 461–470.

50. R. Courtemanche, N. Fujii, A. M. Graybiel, Synchronous, Focally Modulated β-Band Oscillations Characterize Local Field Potential Activity in the Striatum of Awake Behaving Monkeys. J. Neurosci. 23, 11741–11752 (2003).

51. M.-P. Stenner, S. Dürschmid, R. B. Rutledge, T. Zaehle, F. C. Schmitt, J. Kaufmann, J. Voges, H.-J. Heinze, R. J. Dolan, M. Ariel Schoenfeld, Perimovement decrease of alpha/beta oscillations in the human nucleus accumbens. J Neurophysiol. 116, 1663–1672 (2016).

52. R. L. M. Faull, W. J. H. Nauta, V. B. Domesick, The visual cortico-striato-nigral pathway in the rat. Neuroscience. 19, 1119–1132 (1986).

53. E. A. Buffalo, P. Fries, R. Landman, T. J. Buschman, R. Desimone, Laminar differences in gamma and alpha coherence in the ventral stream. Proc. Natl. Acad. Sci. U. S. A. 108, 11262–11267 (2011).

54. G. E. Alexander, M. R. Delong, P. L. Strick, Parallel Organization of Functionally Segregated Circuits Linking Basal Ganglia and Cortex. Annu. Rev. Neurosci. 9, 357–381 (1986).

55. A. D. Smith, J. P. Bolam, The neural network of the basal ganglia is revealed by the study of synaptic connections of identified neurones. TINS. 13, 259–265 (1990).

56. A. M. Graybiel, Building action repertoires: memory and learning functions of the basal ganglia. Curr. Opin. Neurobiol. 5, 733–741 (1995).

57. K. Kunishio, S. N. Haber, Primate cingulostriatal projection: Limbic striatal versus sensorimotor striatal input. J. Comp. Neurol. 350, 337–356 (1994).

58. A. M. Thierry, Y. Gioanni, E. Dégénétais, J. Glowinski, Hippocampo-prefrontal cortex pathway: Anatomical and electrophysiological characteristics. Hippocampus. 10, 411–419 (2000).

