## Supplementary Items for "Volitional Control of Individual Neurons in the Human Brain"

**This PDF file includes:**

Materials and Methods
Table S1
Figs. S1 to S4
References

### Materials and Methods

Experimental Model and Subject Details

Eleven subjects (4 female) aged 28.5±7 years participated in this study (Supplementary Table S1). All subjects were implanted with ADTech Behke Fried Macro-Micro electrodes for the presurgical evaluation of their epilepsy. Implant locations were chosen solely based on clinical indicators. All subjects volunteered for the implantation of microwires and for participation in behavioral tasks, after obtaining informed consent. This study has been approved by the Research Ethics Board of the University Health Network.

Electrode Localization and Electrophysiology

Subjects were implanted with commercially available depth electrodes (Behnke-Fried Macro Micro, ADTech Inc., Racine, MN) which have 8 to 10 macro electrodes along the electrode shaft, and a bundle of 8 microwires spraying out from the tip, plus one ground/reference microwire. The number and location of the electrodes varied from subject-to-subject and were determined based solely on clinical hypothesis of the epileptogenic zones(s). Electrode localization was performed by co-registering pre-op MRI with post-op CT using the iELVIS toolbox (*1*). Following localization, the precise location of each of the electrodes was determined, and later verified by a neurosurgeon.

Electrodes were connected to the Neuralynx Atlas Data Acquisition System (Neuralynx Inc, Bozeman, MT). The macroelectrodes were sampled at 4kHz, with a 16 bit resolution, and bandpass filtered in hardware between 0.1 and 1kHz. The microelectrodes were sampled at 32kHz, with a 16 bit resolution and were bandpass filtered in hardware between 0.1 and 8kHz. A 4-contact subgaleal electrode was used for ground and reference and was placed over the parietal midline facing away from the brain. The microelectrodes were referenced locally, to one of the 8 wires within the same bundle. Neural data was synchronized with behavioural data using TTL triggers sent over Neuralynx's NETCOM protocol.

Online and Offline Spike Detection and Sorting

Online spike detection/sorting was used to drive the neurofeedback task (see *Neurofeedback Task*). Microelectrode channels were bandpass filtered between 300 and 3000 Hz (*2*), and thresholded at 5 times the RMS amplitude. Channels in non-motor regions with well isolated spikes were sorted using the KlustaKwik toolbox in SpikeSort3D (Neuralynx Inc., Bozeman, MT). Sorted templates for each channel were then sent back to the recording software, Pegasus (Neuralynx Inc., Bozeman MT). The instantaneous timestamps of sorted neurons were then streamed to custom written scripts in MATLAB (Mathworks Inc., Natick, MA) over the Neuralynx NETCOM protocol. Although online spike sorting was performed on numerous channels, only the online spike train of the direct neuron (i.e. the neuron being trained) was used for post-hoc analysis. The remainder of the neurons (i.e. the indirect neurons) were detected and sorted offline.

For offline spike detection, all microelectrode channels were bandpass filtered between 300-3000Hz, and spikes were subsequently detected using threshold crossings of a local energy measurement, calculated by convolving the raw signal with a kernel of approximate width of an action potential. All detected spikes were sorted using the semiautomatic template-matching algorithm OSort, that is available as open source (Rutishauser et al., 2006). Similar to previous work (*3*), we classified clusters as putative single neurons by looking at the following criteria (1) violation of refractory period, (2) shape of the ISI distribution, (3) shape of the waveform, and (4) separation from other clusters. Clusters that appeared similar to one another were merged, and clusters that were either contaminated with noise, or failed the meet the criterion described above were rejected. For the accepted clusters, the individual waveforms, along with the timestamp of each spike and its cluster definition were saved.

Neurofeedback Task

We developed a novel, low-latency intracranial neurofeedback task in which the vertical movement of a red square on a screen was controlled by the smoothed instantaneous firing rate of a well isolated neuron (called the direct neuron) from a mnemonic, non-motor structure in the human brain (Table S1). The direct neuron was chosen by the experimenter such that it was (1) well isolated from background noise, (2) not contaminated by movement artifacts, (3) had less than 3% of inter spike intervals below 3ms, and (4) had a baseline firing rate greater than 0.5Hz. . The spike timings of the chosen, online sorted direct neuron were streamed to custom scripts in MATLAB using the Neuralynx NETCOM protocol. The spike timings were used to create a spike train, which was stored in a 2-second first-in-first-out buffer, which was updated every ~40 ms. This spike train was then smoothed by convolving it with a 200ms Gaussian kernel. The instantaneous value of the smoothed spike train was used as the control signal. Our visual neurofeedback task was developed using Psychtoolbox, and consisted primarily of a red square capable of moving vertically on a screen, along with a white horizontal line indicating the target (Figure 1). The vertical movement of the red square was controlled by the described control signal. Each participant was asked to modulate their brain activity to move the square above the target line and hold it there for at least 0.5 seconds. Every time the block went above the target line, its colour was changed to purple to clearly identify to the participant that the block was above the line.

Each testing session began with a ~4 minute baseline session in which participants stared at a fixation cross for 1 minute, followed by a 1 minute eyes-closed period, followed by passively viewing a red square moving on the screen for 2 minutes. All participants were instructed to minimize body movements throughout the session, and any trials with overt physical movements were noted, and later rejected. Following the baseline session, an appropriate, well-isolated neuron with a firing rate above 0.5Hz was chosen as the neuron to be trained. If the subject participated in more than one training session, we ensured to choose a neuron from a different electrode, in order to ensure independence of each session. Using this well-isolated neuron, we then performed a short, ~5 minute training and familiarization session in which we asked the participant to modulate their brain activity to move the block on the screen. This session was also used to determine an appropriate starting difficulty for the task (i.e. the position of the horizontal target bar on the screen). An appropriate difficulty was defined as a multiple of the standard deviation of the baseline firing rate of the selected neuron, chosen such that the participants required ~60 seconds to achieve success. In order to prevent biasing each participant's unique control strategy, we never provided the participants with explicit instructions on how to modulate their neural activity. Furthermore, literature suggests that participants who report no specific control strategies demonstrate better neurofeedback performance than those who report specific mental strategies (*4, 5*).

Following the familiarization/ training portion, we moved onto the primary testing portion of the session. This portion was divided into several blocks, consisting of 10 trials each.

Each trial began with a start message, following by a fixation cross (3 seconds), followed by the actual trial in which the participants volitionally moved the red square on the screen. Each trial continued until the participant was successfully able to push the block above the line and hold it there for 0.5 seconds. This resulted in a "Trial Successful" message (1 second) followed by a distractor math question (i.e. addition of 3 integers between 1 and 5). Answering the question triggered the start of the next trial. Participants were instructed to try and complete each block of trials (i.e. 10 trials) in 10 minutes or less. If the participants successfully completed all 10 trials in less than 10 minutes, the difficulty of the next trial was marginally increased (by increasing the target threshold by 0.2 standard deviations). Participants were required to complete at least 3 blocks of testing, or 30 trials. Sessions in which testing was interrupted before the minimum number of trials were completed were not used in the analysis. Testing was continued up to a maximum of 12 blocks, or stopped earlier if otherwise interrupted by visitors, clinical interventions or self-reported fatigue. After the testing session, a post-testing baseline session was performed, which mirrored the pre-testing baseline session described above.

### Data Analysis

All analyses were performed in MATLAB with custom written routines and with the aid of open-source toolboxes where possible. All LFP data was low-pass filtered at 250Hz, downsampled to 1kHz and notch filtered at 60, 120, 180 and 240Hz. Online and offline spike times were used to create binary spike trains with a bin width of 1ms (corresponding to 1kHz sampling). Firing rates were calculated by convolving the binary spike trains with a scaled 200ms gaussian. Average firing rates correspond to the mean of the smoothed spike train in each trial. Peak firing rate corresponds to the maximum of the gaussian convolved smoothed spike train in each trial. Modulation depth (MD) of each neuron was calculated as the average firing rate in the 1 second window before success minus the average firing rate in the 1 second window after success. Bursts were detected using a non-parametric version of the commonly used Poission-Surprise method, called Rank-Surprise method (*6*), which identifies bursts based on the probability that a given number of spikes occur in a given duration, given the overall distribution of the inter-spike-intervals for a particular neuron. Burst frequency was defined as the total number of bursts in each trial divided by the duration of each trial. A session was marked as "Learner" if a linear regression of the average or peak firing rate with the trial number resulted in a significant positive slope. i.e. "Learner" sessions were those in which there was a significantly positive trend in the average and/or peak firing rates within a single testing session. All other sessions were marked as "Non-Learner". In all analyses, the term "early" corresponds to the data from the first 15 trials of the testing session, and "late" corresponds to data from the last 15 trials of the testing session. This definition was chosen since the minimum number of trials completed by each participant was 30.

All spectral analyses were performed using the open-source Chronux toolbox (http://chronux.org) for MATLAB. Spectral estimates were obtained using a multitaper method, with a total of 5 tapers and a time-bandwidth product of 3. For all spectrograms, a moving window size of 1 second was used (to accurately resolve frequencies as low as 1Hz), with a step size of 50ms. Spike field coherence was calculated as:

$$C = \frac{|R_{xy}|}{\sqrt{|R_{xx}|}\sqrt{|R_{yy}|}}$$

where $R_{xx}$ and $R_{yy}$ are the power spectra of the spike train and LFP oscillation respectively, and $R_{xy}$ is the cross-spectrum. All local spike-field coherence (SFC) values were calculated between any given putative neuron and the closest macro electrode to ensure that the spiking activity itself did not contaminate the power spectrum of the recorded oscillations. Since SFC estimates can be affected by the firing rate of the selected neuron, we performed a probabilistic spike thinning procedure in order to equate the firing rate of the early and late trials (*7*). To do so, we convolved the binary spike trains with a 10ms Gaussian kernel, and averaged them across trials. Next, we determined the probability that a spike should be removed at any given point in time by subtracting the early and late firing rates at each point in time, and dividing by the maximum rate in the early and late trials at that point in time. Using this probability train, we randomly removed spikes from the original spike trains. For example, if the probability value at any given point in time was 50%, then the probability that a spike in the late trials at that point in time was removed was 50%, resulting in roughly 50% of the spikes at that point in time being removed. We verified that the spike thinning procedure successfully eliminated any differences between the firing rate between the early and late trials (Supplementary Figure S2).

Quantification and Statistical Analysis

Unless otherwise specified, the bold line in all figures corresponds to the mean of the data, and the shaded area and/or the error bars correspond to the SEM. In all box plots, the shaded box corresponds to the SEM, and the bold vertical line corresponds to the standard deviation. All means, standard errors and statistical tests were calculated across sessions. The number of subjects or sessions contributing to each figure is indicated with the corresponding N value. Wherever necessary, data outliers were removed using the Grubbs outlier test. Wherever possible (given data normality and absence of outliers), parametric tests were used to test for significance. Otherwise, non-parametric equivalents were used. When determining significant fluctuations in averaged waveforms (time-aligned or power spectra), a non-parametric permutation test was using with random time shuffling, and 2000 iterations. The specific statistical test used for each figure is stated clearly in the text and/or in the figure legend. The data epoch being analyzed in any particular figure is cleared labelled and described in the figure legend. Significance for all statistical tests was set at $p<0.05$. A single asterisk indicates significance at the $p<0.05$ level, double asterisk indicates significance at the $p<0.01$ level and a triple asterisk at the $p<0.001$ level. All statistical tests were performed using either custom scripts or built-in functions in MATLAB.

**Table S1 – Participant Demographics**

| Session ID | Patient ID | Sex | Handedness | Learner Session? | Direct Neuron Anatomy | # Trials Completed | Age | Notes |
|---|---|---|---|---|---|---|---|---|
| 1 | 1 | M | R | Y | Lateral Orbitofrontal | 50 | 25.1 | |
| 2 | 1 | M | R | Y | Lateral Orbitofrontal | 53 | 25.1 | Missing Macro Data |
| 3 | 2 | F | R | N | Amygdala | 39 | 46.1 | |
| 4 | 3 | F | R | Y | Hippocampus | 117 | 20 | |
| 5 | 3 | F | R | N | Insula | 77 | 20 | |
| 6 | 4 | F | L | Y | Hippocampus | 77 | 25.5 | |
| 7 | 5 | M | R | Y | Amygdala | 30 | 26.2 | |
| 8 | 5 | M | R | Y | Amygdala | 39 | 26.2 | |
| 9 | 6 | M | R | Y | Lingual Cortex | 59 | 38.1 | |
| 10 | 6 | M | R | Y | Hippocampus | 59 | 38.1 | |
| 11 | 7 | M | R | Y | Hippocampus | 94 | 28.5 | |
| 12 | 8 | F | R | N | Hippocampus | 48 | 25.9 | Missing Macro Data |
| 13 | 9 | M | R | N | Hippocampus | 48 | 24.6 | |
| 14 | 9 | M | R | N | Amygdala | 52 | 24.6 | |
| 15 | 10 | M | R/L | N | Hippocampus | 29 | 25.3 | |
| 16 | 10 | M | R/L | Y | Hippocampus | 48 | 25.3 | |
| 17 | 11 | M | R | N | Amygdala | 47 | 28.1 | |

**Figure S1 – Brain-Wide indirect neurons are also not task-contingent**

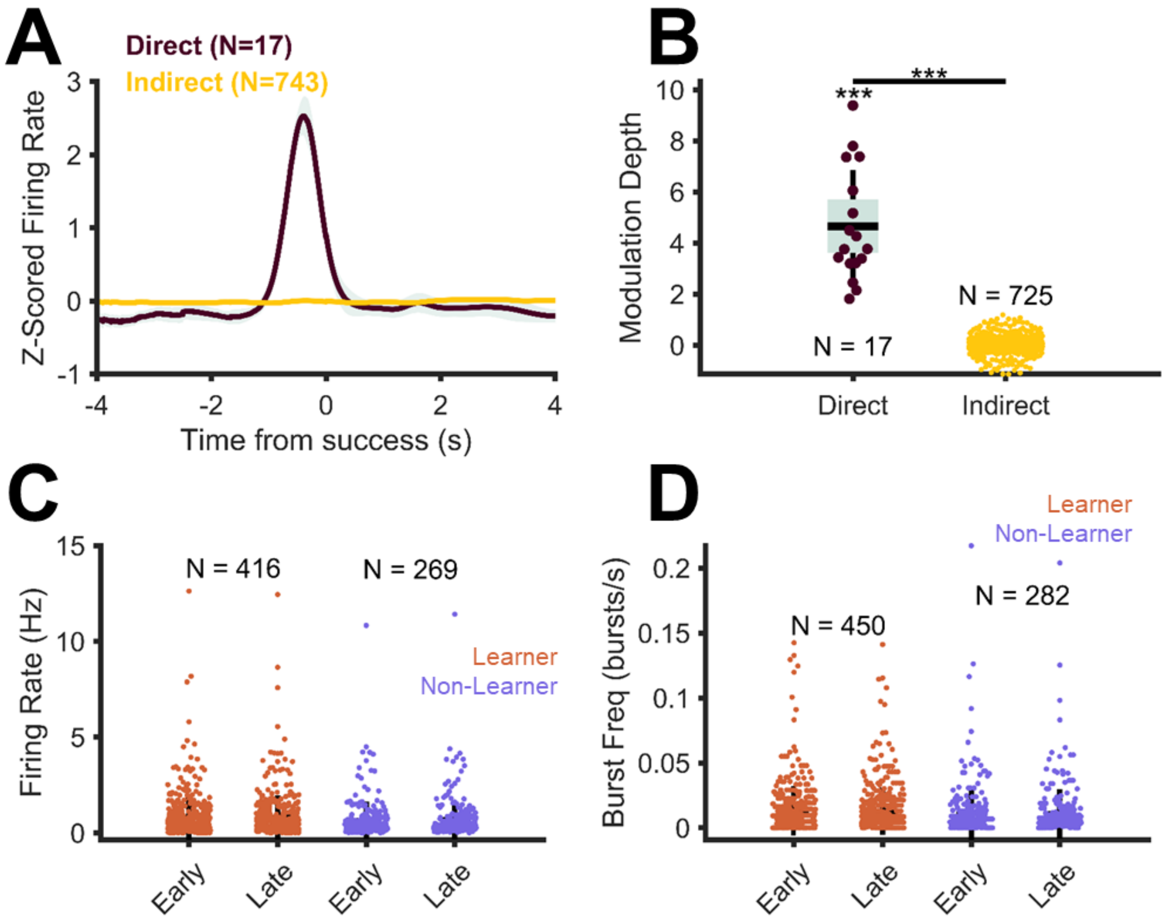


(A) Firing rate of the direct neuron increases sharply immediately before success. Firing rate of all other recorded neurons (from all other microwire bundles) is not modulated around success.
(B) Modulation depth of direct neurons is significantly greater than zero ($p<0.001$, single sample two-tailed T Test) and it is significantly greater than that of brain-wide indirect neurons ($p<0.001$, independent samples T Test). Outliers are removed using the Grubbs method.
(C) Changes in the firing rate of brain-wide indirect neurons within a single session (Early trials = first 15 trials, late trials = last 15 trials) grouped by learner and non-learner sessions. Firing rate of the brain-wide indirect neurons does not change for the learner or non-learner sessions.
(D) Same as (C) but for burst frequency. Burst frequency of the indirect neurons also does not change for either the learners or the non-learners.

**Figure S2 – Probabilistic Spike Thinning Diminishes early vs. late differences in firing rate of direct neurons.**

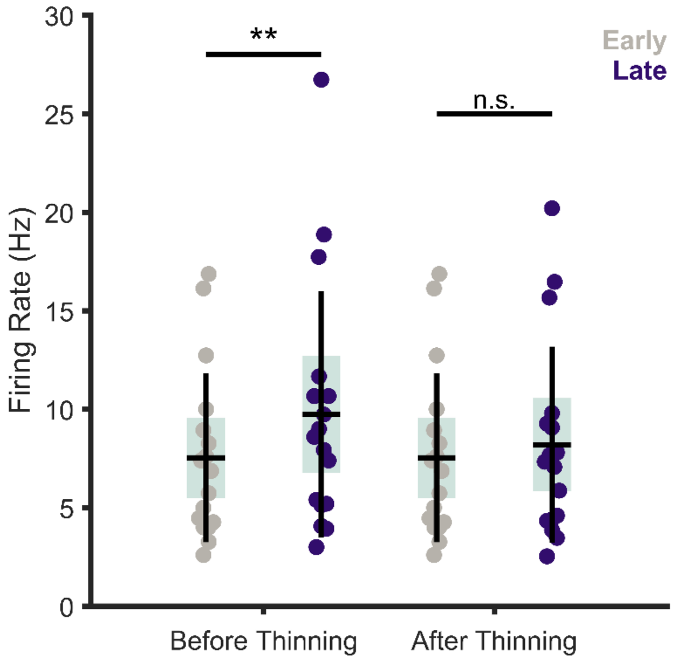


Results of the spike thinning procedure. The firing rate in the early versus late trials is significantly different before the spike thinning procedure ($p = 0.0052$, Paired T Test). However, after performing the spike thinning procedure, the firing rate is not different between the early and late trials ($p = 0.11$, Paired T Test).

**Figure S3 – Learning Related Changes in Spike-Field Coherence between Direct Neurons and Non-Local LFP.**

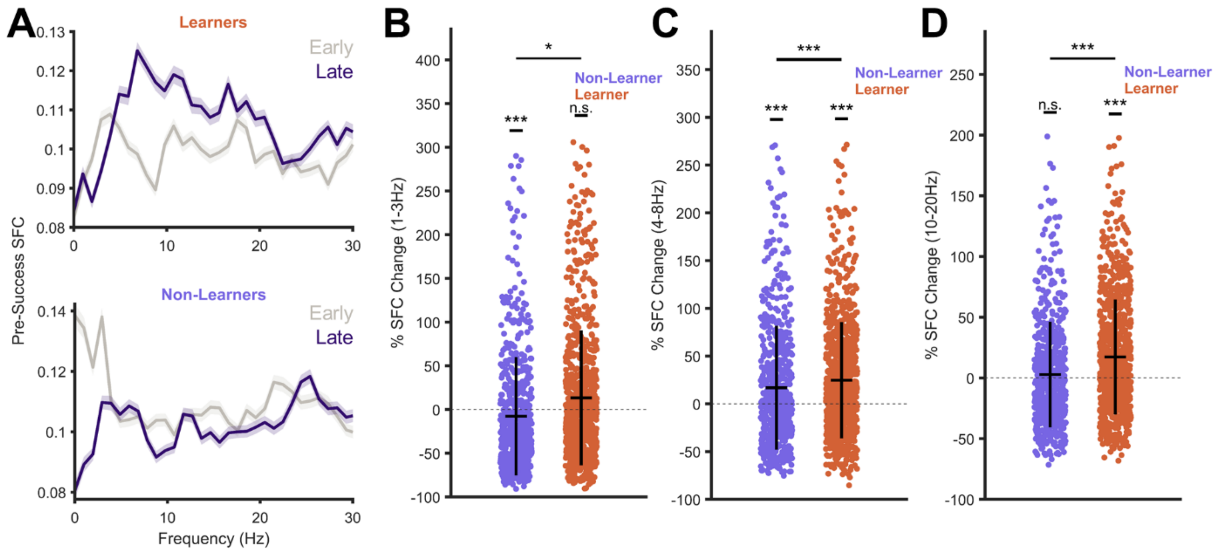


(A) Change in SFC between direct neurons and non-local LFP recorded at macro electrodes throughout the brain, for early and late trials, separated for learner (top) and non-learner (bottom) sessions. Note that the change in SFC between early and late trials for the learners appears to have a peak at 6Hz, contrary to the peak at ~12Hz for the SFC calculated between the direct neurons and the local LFP (Figure 3).
(B) Percent (%) change in SFC in the delta (1-3Hz) frequency band from early to late trials for learners and non-learners. SFC in the delta band decreases significantly in the non-learner population ($p=1.06*10^{-7}$, Wilcoxon Sign Rank Test), but not in the learners ($p = 0.57$, Wilcoxon Sign Rank Test). There is also a significant difference in the SFC change between the learner and non-learner populations ($p = 0.021$, Wilcoxon Rank Sum Test).
(C) Same as B, but for the theta (4-8Hz) frequency band. SFC in the theta band increases significantly in the learner and non-learner sessions ($p = 3.3*10^{-21}$ for learners, $p = 4.2*10^{-4}$ for non-learners, Wilcoxon Sign Rank Tests). The increase is significantly greater for the learners than the non-learners ($p = 6.1*10^{-12}$, Wilcoxon Rank Sum Test).
(D) Same as B and C, but for the 10-20Hz frequency band. SFC in the learners increases significantly between the early and late trials ($p = 5.8*10^{-16}$, Wilcoxon Sign Rank Test), but not for the non-learners ($p = 0.99$, Wilcoxon Sign Rank Test). The increase in learners is also significantly greater than the change in the non-learners ($p = 9.4*10^{-10}$, Wilcoxon Rank Sum Test). For all panels, * represents $p<0.05$, *** represents $p<0.001$. All tests are corrected for multiple comparisons using the Bonferroni method.

**Figure S4 – Changes in spectral power following unsuccessful threshold crossing**

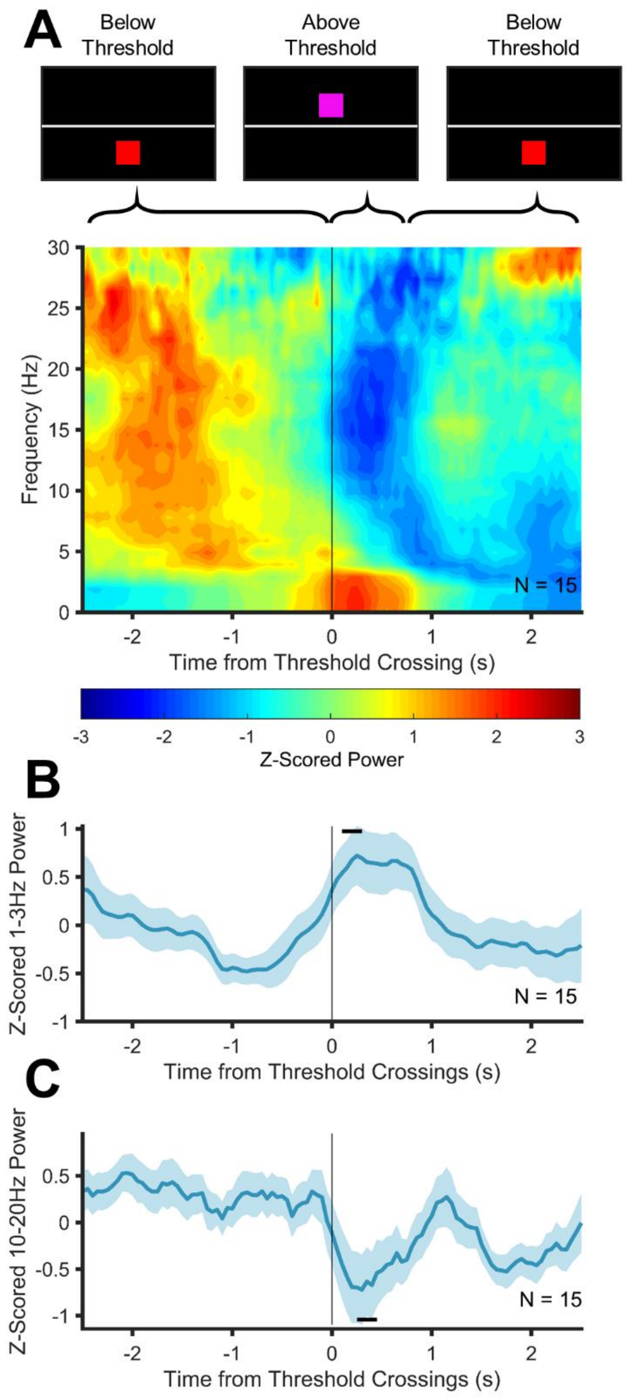


(A) Normalized power spectrogram aligned to threshold crossings, averaged across all subjects. Note significant increase in delta power immediately following threshold crossing.
(B) Grand average Z-Scored delta power (1-3Hz) aligned to threshold crossings. Note that the power increases significantly immediately following threshold crossings.
(C)Same as (B) but for the 10-20Hz power. Notice the significant decrease in 10-20Hz power immediately following threshold crossings.
For (B) and (C), significant time-points are marked with a bold black line. Significance tested using a non-parametric, random-shuffling permutation test with. Significance level set to $p<0.05$. Shaded areas represent the SEM.

**References**

1. D. M. Groppe, S. Bickel, A. R. Dykstra, X. Wang, P. Mégevand, M. R. Mercier, F. A. Lado, A. D. Mehta, C. J. Honey, iELVis: An open source MATLAB toolbox for localizing and visualizing human intracranial electrode data. *J. Neurosci. Methods*. **281**, 40–48 (2017).
2. U. Rutishauser, E. M. Schuman, A. N. Mamelak, Online detection and sorting of extracellularly recorded action potentials in human medial temporal lobe recordings, in vivo. *J. Neurosci. Methods*. **154**, 204–224 (2006).
3. M. C. M. Faraut, A. A. Carlson, S. Sullivan, O. Tudusciuc, I. Ross, C. M. Reed, J. M. Chung, A. N. Mamelak, U. Rutishauser, Data Descriptor: Dataset of human medial temporal lobe single neuron activity during declarative memory encoding and recognition (2018), doi:10.1038/sdata.2018.10.
4. S. E. Kober, M. Witte, M. Ninaus, C. Neuper, G. Wood, Learning to modulate one's own brain activity: The effect of spontaneous mental strategies. *Front. Hum. Neurosci.* **7**, 1–12 (2013).
5. R. Sitaram, T. Ros, L. Stoeckel, S. Haller, F. Scharnowski, J. Lewis-Peacock, N. Weiskopf, M. L. Blefari, M. Rana, E. Oblak, N. Birbaumer, J. Sulzer, Closed-loop brain training: The science of neurofeedback. *Nat. Rev. Neurosci.* **18** (2017), pp. 86–100.
6. B. Gourévitch, J. J. Eggermont, A nonparametric approach for detection of bursts in spike trains. *J. Neurosci. Methods*. **160**, 349–358 (2007).
7. G. G. Gregoriou, S. J. Gotts, H. Zhou, R. Desimone, High-Frequency, long-range coupling between prefrontal and visual cortex during attention. *Science (80-. ).* **324**, 1207–1210 (2009).
